# Parameter Identifiability and Non-Uniqueness In Connectome Based Neural Mass Models

**DOI:** 10.1101/480012

**Authors:** X. Xie, A. Kuceyeski, S.A. Shah, N.D. Schiff, S.S. Nagarajan, A. Raj

## Abstract

The spatial-temporal patterns of neuronal dynamics emerge from the network of coordinated brain regions, this structure-function relationship of the brain can be described mathematically by biophysical models of coupled brain regions connected by white matter tractography. Implementations of such models have focused on reproducing functional connectivity extracted from functional magnetic resonance imaging (fMRI), but these efforts are limited by the temporal resolution of fMRI data and the reduction of time course recordings into phenomenological functional connectivity maps. Here, we optimize parameters of a neural mass model (NMM) to best fit region-wise power spectra across the whole brain estimated from source localized electroencephalography (EEG). NMM models with global parameters were not able to fully reproduce region-wise power spectra, with or without the inclusion of structural connectivity information. In contrast, without the inclusion of structural connectivity information, independent oscillators at each brain region are able to reproduce region-wise power spectra. But the addition of structural connectivity and transmission delays to the NMM does not improve overall power spectra fit. Connectome-based NMM implementations with regional parameters lead to high dimensional network models that produce non-unique results. Inherent parameter identifiability problem in network models poses challenges for using such models as diagnostic tools for neurological diseases.

## 1. Introduction

Neuronal populations in the brain are continuously active whether they are engaged in behavior or at rest, and it is evident that functional activity produced by interacting neuronal populations is a product of specific neuronal properties as well as connection strength amongst themselves [32, 44, 7, 40]. One powerful approach to mathematically describe this structure-function relationship is the simulation of neural mass models (NMMs). This approach models the average activity with a small number of parameters to summarize the behavior of a neural ensemble [31, 39, 56, 8]. A neural ensemble is a set of locally interacting neurons [16], and the properties of these neurons can be described in terms of their mean firing rate and mean membrane potentials, therefore a NMM can represent the lumped activity of a specific neuronal cell type or a particular functional brain region [49, 50]. By coupling several such lumped neural masses via structural connectivity and transmission delay, these models are capable of producing functional connectivity in both healthy[18, 32, 5] and disease[53, 6, 51].

Previous macroscopic network models of brain dynamics relied on the spatial resolution of fMRI signals [4, 17] or encephalography data [11, 14] in source space to identify functionally related brain regions and evaluated model performance based on band-limited functional connectivity (FC) metrics. FC metrics are a statistical map of correlated interactions between brain areas, this phenomenological metric is not sufficient to fully explain the temporal dynamics between brain regions[9]. Thus, theoretical models limited to reproducing selective FC metrics are under-utilized and do not fully address the structure-function relationship of the brain. However, fMRI data is limited by its temporal resolution and can only contain signals in the low frequency bands, does not capture neural oscillatory frequencies observable in electrophysiological recordings like electrocorticography (ECoG), magnetoencephalography (MEG), and EEG.

The observable alpha, beta, gamma, delta and theta rhythms follow a spatially distributed pattern [3, 22, 33, 37, 35]. For example, the alpha range is distinctively shown in the occipital lobe and posterior temporal cortex [25, 46, 41, 43]. These heterogeneous patterns of the observable power spectra are produced by a combination of local oscillatory behavior at brain regions and varying degrees of connectivity between distant brain regions. Previous works in modeling structure-function relationships have mostly assumed the same local properties across the network to reproduce fMRI functional connectivity patterns [9, 20, 57], but exploring the frequency profiles for each brain region using such a simplistic approach has not been explored. While it is encouraging that oscillator models are capable of displaying expected frequency behavior [8, 49, 50, 54] and can reproduce functional connectivity to a limited extent [10, 45, 47], the addition of SC and transmission delay to local parameters in a network model has not been explored in terms of how well they can predict regional power spectra.

Here, we first implemented a Wilson-Cowan oscillator NMM with or without SC. We estimated the source localized EEG power spectra and then optimized model parameters of variants of our NMM to fit the regional power spectra estimates. We implement three variations of the Wilson-Cowan Oscillator model as illustrated in Figure 1; 1) varying oscillators (VO) at each brain region without structural connectivity to check if the model is able to simulate power spectra across all brain regions without SC; 2) identical oscillators at each brain region with structural connectivity (IOC) to observe model fitting with SC but without any variations in local circuitry; 3) varying oscillators at each brain region with structural connectivity (VOC) to fit empirical source localized EEG power spectra with both SC and local circuitry variation. The resulting optimized models show local parameter variations is needed to generate spatially varying patterns of power spectra, furthermore, the addition of SC and global parameters in higher dimensional models did not improve model fitting to observed EEG spectra.

**Figure 1:**
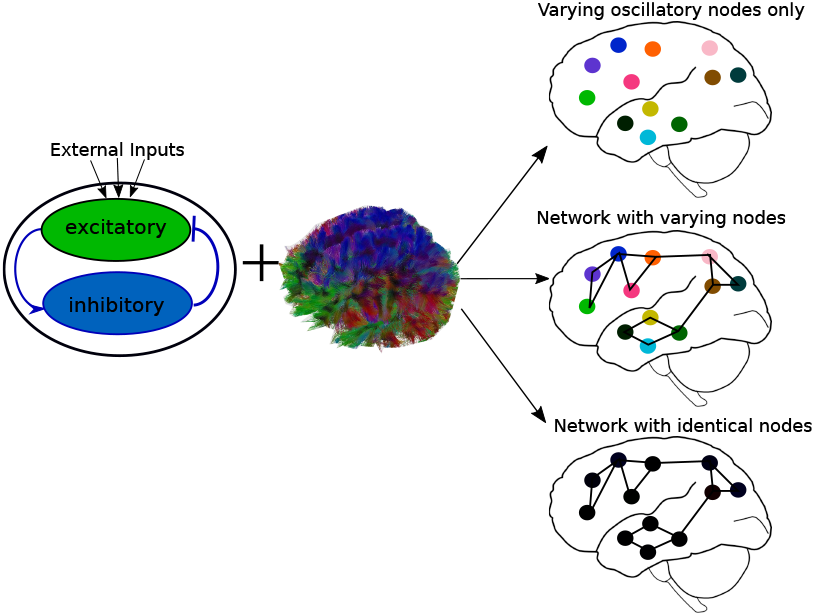
Variations of the Wilson-Cowan model. Varying oscillators (VO) at each node without connectivity, varying oscillators at each node plus connectivity (VOC), or identical oscillators at each node plus connectivity (IOC)

These high dimensional models require inference of a high number of parameters. Accurate inference of the model parameters in a complex network of interacting brain regions is incredibly difficult for any optimization method. Consequently, the over-specification of such models may result in non-unique solutions. Indeed, we find that higher dimensional connectome coupled NMMs are not able to provide uniquely identifiable solutions to the parameter inference problem. This poses a challenge for the emerging idea of using inferred model parameters as diagnostic biomarkers of neurological diseases[43, 13, 26, 48].

## 2. Methods

### 2.1. Subjects and Data Collection

All experiments were conducted after obtaining written informed consent from the subjects and approval by The Institutional Review Board of Weill Cornell Medical College. T1-weighted anatomical MRI and diffusion-MRI scans were collected from 11 out of the 13 healthy individuals (8 male, 35.2 +/- 12.25 years) on a 3.0 Tesla General Electric Signa Excite HDx (GE Healthcare, Waukesha, WI) clinical MRI system with an eight-channel head receive-only coil. DMRI scans were obtained using a spin-echo diffusion tensor pulse sequence with one T2-weighted image, 33 diffusion-weighted images (one subject is an exception with 55 directions) evenly distributed on a sphere with b = 1000 s/mm2, TE = 76.7 ms, TR = 9000 ms, field of view = 22 cm, 28 slices of 5.0 mm thickness, matrix size = 128 × 128, reconstructed with zero filling to 256 × 256. An axial 3D IR-prepped, fast SPGR with parameters tuned to optimize brain tissue contrast sequence (BRAVO sequence) was used for anatomical imaging with inversion time = 400 ms, TR = 8.9 ms, TE = 3.5 ms, flip angle = 13 degrees, axial field of view = 24 cm, 136 slices of 1.2 mm thickness, matrix size = 256 × 256, parallel imaging acceleration factor = 2. Additionally, eyes-open (EO) and eyes-closed (EC) Resting-state EEG data was collected for 9 out of the 13 healthy subjects. Recordings for a minimum of 110 seconds were performed with a 129-channel HydroCel Geodesic EEG Sensor Net (Electrical Geodesics, Eugene, Oregon). The impedance of all electrodes was < 75*k*Ω at the beginning of the recording, the EEG signals were sampled at 250 Hz sampling frequency and filtered from DC to 100 Hz. Datasets were chosen for analysis only if all data modalities were present without unacceptable levels of noise or artifacts. Only 7 subjects had complete sets of usable EEG, MRI, and DTI data, so we proceeded with analyses using only those subjects.

### 2.2. Structural Connectivity Networks

Structural and diffusion MR volumes were co-registered and pre-processed in the manner previously described [29]. Segmentation of gray matter, white matter, and cerebrospinal fluid was performed after slice-timing correction, realignment, co-registration and/or normalization, and spatial smoothing was performed using SPM8 (Statistical Parametric Mapping tool). The gray matter was further parcellated into 86 anatomical regions of interest (ROIs) based on the Desikan-Killany atlas using the established FreeSurfer package [15]. The parcellated regions were used to seed tractography nodes in coregistered diffusion MRI volumes. The connectivity between any two regions was given by a weighted sum of tracts going between them as described by [23]. The algorithm traces likely white matter fiber tracts by taking into account tissue probability maps as well as diffusion orientation in a Bayesian manner, the tracing stopped when the track angle between steps exceeded *pi/*3 or when encountering a voxel that is outside of the white matter mask.

### 2.3. Source Localization

Source localization of the EEG signals was performed with Brainstorm [52], which is documented and freely available for download online under the GNU general public license (http://neuroimage.usc.edu/brainstorm). Prior to source localization, the raw EEG data were band-pass filtered between 2 and 45 Hz, transience time segments and unusable channels were manually removed after inspecting the time series and its power spectrum. We then applied an average reference followed by independent component analysis to remove artifacts such as eye blinks and heart beats that are picked up by the EEG electrodes, removal of additional noisy time segments was performed manually after inspection.

Source localization was performed with a “warped” Colin27 template head model to remove variations due to noise level, head position, and starting/ending slices for MRI acquisition runs. The Colin27 template is a stereotaxic average of 27 T1-weighted MRI scans of a single individuals head [19]. To incorporate individual subject’s anatomical information, we created pseudo-individual anatomies using Brainstorm’s warp anatomy functions to deform and scale the high resolution Colin27 head shapes to match each subject’s individual head shapes. Surface meshes of the brain, skull, and scalp were extracted from the template MRIs using 1922 vertices per layer. To obtain an analytical approximation of the lead field for the conductive brain volume, we chose to use the three-shell spherical harmonics expansion methods as discussed by [38]. Specifically, an initial grid of 4000 source points was generated from the cortex surface and samples the brain volume in an adaptive manner towards the center of the brain, each grid layer is down-sampled by a factor of 3 for a maximum of 17 layers, resulting in a total of 11151 to 16442 dipole sources depending on individual head anatomy. A representative visualization of the dipole sources is shown in Fig. S1.

To obtain the inverse solution, a noise covariance matrix was calculated over the EEG recordings to model the noise contaminating our data; only the diagonal elements were kept for the inverse solution to estimate the variance of each sensor. For all subjects, the activity at each dipole source was estimated using a linearly constrained minimum variance (LCMV) spatial filter [55]. Three-dimensional dipole sources yielded a 4D time series (*x* × *y* × *z* × *time*) for each set of EEG recordings. The norm of the 3 spatial coordinates 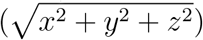 at each time point was taken to produce a 1D time series of estimated activation over the entire dipole. An average time series was obtained for all sources belonging to each of the same 86 ROIs as defined previously (See Fig. S1 for visualization of the dipoles), and the source localized time series were used as empirical data for modeling training.

### 2.4. Wilson & Cowan Neural Mass Model

To model neurophysiological activity from anatomical architecture for each ROI, we adopt the Wilson-Cowan coupled oscillators [56]. This model assumes that a local circuit consists of two lumped masses of excitatory and inhibitory neural populations interacting with each other, whole brain regional dynamics are achieved by coupling local masses via structural connectivity *A_jk_*, global coupling parameter *c*_5_, and a transmission delay 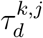.

The simulated average activity at the *jth* brain region is:

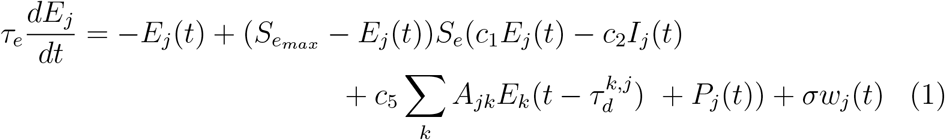

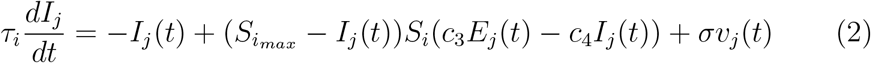

Where *E*(*t*) and *I*(*t*) represent the firing rate of the excitatory and inhibitory neuronal populations respectively, *τ* is a time constant and *w_j_*(*t*) and *v_j_*(*t*) are random normally distributed noise with standard deviation *σ*. *P*(*t*) is an external input parameter to the excitatory neural ensemble that controls oscillatory activity, local parameters *c*_1_, *c*_2_, *c*_3_, and *c*_4_ represent the average number of excitatory and inhibitory synapses within a neuronal ensemble. *S_e_* and *S_i_* are transfer functions characterized by the sigmoidal function capturing the non-linear response of a cell generating an action potential based on summed synaptic input:

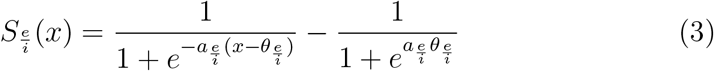

Different variations of this model (Fig. 1) can simulate average neuronal activity at each region in the brain. Here, we will compare three models (1) the varying oscillator (VO) model that consists of varying local neuronal ensemble with only locally defined parameters and no inter-connectivity between nodes, (2) the varying oscillator plus connectivity (VOC) model that consists of local neuronal ensembles with varying local parameters, plus a global coupling parameter, structural connectivity, and transmission delay, and (3) the identical oscillators plus connectivity (IOC) model that consists of local neuronal ensembles with uniform local parameters, plus a global coupling parameter, structural connectivity, and transmission delay.

### 2.5. Evaluating Oscillatory Abilities of the Neural Mass Model

To assess if the neural mass models are able to produce a frequency profile that covers all signature physiological frequency bands, we performed 2-seconds simulations with varying parameters. Firstly, simulations at a single node with no connectivity were performed with varying excitatory and inhibitory time constant parameters (*τ_e_, τ_i_*) operating in the range 1*ms*−40 *ms* with a step size of 1 *ms* and an external driving parameter of *P*(*t*) = 2.5. When the structural connectivity matrix is introduced, the global coupling parameter *c*_5_ and transmission velocity also dictate oscillatory activity. For the 86-region network model, we varied the global coupling parameter from 0 to 3 with a step size of 0.2. Upon identifying the value of *c*_5_ for which the network model transitioned to oscillatory behavior (as done previously in [36]), additional 1-second simulations were performed with varying transmission velocity from 5 *m/s* to 50 *m/s* with a step size of 5 *m/s*. The power spectra of each simulation were computed to select the peak oscillatory frequency. All power spectra calculations were performed with MATLAB’s multi-taper power spectral density estimate function *PMTM*. Simulations were performed with default local parameters as illustrated in [36]: *c*_1_ = 16*, c*_2_ = 12*, c*_3_ = 15, *c*_4_ = 3, and sigmoidal function parameters: *a_e_* = 1.3, *a_i_* = 2, *θ_e_* = 4, *θ_i_* = 3.7.

### 2.6. Model Optimization

The model was implemented using simulation runs of 3 seconds, using a numerical integration time step of Δt = 0.004 sec or 250 *Hz* with MATLAB’s *ode45* function. The noise term in the model was removed to maintain an unchanging parameter space during optimization. To improve the odds that we capture the global minimum of a suitably defined goodness of fit (GOF) criterion in our parameter space, we chose to implement the probabilistic approach of simulated annealing [28]. The algorithm samples a very large set of parameters within a set of boundaries by generating an initial trial point and choosing the next trial point from the current point by a probability distribution with a scale depending on the current “temperature” parameter. While the algorithm always accepts new trial points that map to cost-function values lower than the previous cost-function values, it will also accept trial points that have cost-functions with greater values than the previous point to move out of local minima. The acceptance probability function is 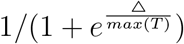, where T is the current temperature and Δ is the difference of the new minus old cost-function values.

Our cost-function was defined as the two-sample Kolmogorov-Smirnov (KS) statistic between the empirical source localized spectra and simulated spectra from each model variation. The initial parameter value and boundary constraints for each parameter are given in Table 1; these had the same values regardless of model variation.

**Table 1:** Initial values and boundary constraints for all model parameters in the simulated annealing optimization

|  | Initial Value | Lower/Upper Boundary |
| --- | --- | --- |
| Time constants $\{\tau_e, \tau_i\}$ | 20ms | [5ms, 30ms] |
| Local Parameters $\{C_1, C_2, C_3, C_4\}$ | 16, 12, 15, 3 respectively | [1, 20] |
| Global Coupling $C_5$ | 1.5 | [0, 10] |
| Transmission Velocity | 10m/s | [5m/s, 30m/s] |
| External Input $P(t)$ | 2.5 | [2.0, 3.0] |

All simulated annealing runs were allowed to iterate over the parameter space for a maximum of *N_p_ ×* 500, where *N_p_* is the number of parameters in the model. To ensure the optimization algorithm thoroughly scanned the parameter space and arrived at a global minimum within the boundary constraints, the initial temperature was raised to 200 (default = 100) for all parameters, and the cooling schedule was set to the average of the quotient between initial temperature and the iteration number for each parameter. Such a cooling schedule ensures that the temperature is low at high iteration counts, so that the optimization algorithm only travels along the downward slope of the current minimum. The VO model was optimized first to obtain parameters for time constants, local parameters, and the external drive parameter. Then these local parameters were fixed in the VOC model optimization that focused on the global parameters of global coupling and transmission velocity. The IOC model’s optimization was performed to identify global parmaeters and one set of local parameters for all 86 brain regions. To ensure that we reached the optimal parameters for the VOC model, we performed an additional optimization where the local parameters were allowed to vary. A conditional minimization algorithm was employed where simulated annealing was performed alternatively for local parameters and global parameters over 10 iterations (VOC-CM). Upon the 10th iteration, four subjects showed slight decreases in cost-function evaluation from the 9th iteration. Upon further inspection, their changes in cost-function was smaller than 0.5% from the previous iteration. To ensure convergence, we continued their optimization to 15 iterations to avoid local minima.

### 2.7. Model Performance and Analysis of Simulated Power Distribution

Simulated power spectra were obtained after reintroducing the Gaussian noise term (*σ* = 0.00001) back into the model and allowing it to run for the duration of the simulations. We calculated the average spectra over 10 different model simulations to account for noise for each set of optimized parameters. Each brain region’s source localized and simulated power spectra were split into alpha (8 − 12 *Hz*) and beta (12 − 25 *Hz*) bands, the total power in each band were computed by summing the normalized power after subtracting the mean at each frequency bin. Visualization of regional alpha and beta band power are displayed on glassbrains generated with an opensource tool “Brainography” developed by our group [30].

We also computed the Kolmogorov-Smirnov statistic between the source localized spectra and each model variations simulated spectra for each brain region. Due to the non-Gaussian distribution in the Kolmogorov-Smirnov statistic at the end of all simulations, a Wilcoxon rank-sum test was used to compare the distribution of Kolmogoriv-Smirnov statistics between the three model versions. All parameters that fell within ± 1% of the median optimized Kolmogorov-Smirnov statistics in VO and VOC-CM were extracted for visualization of their distribution.

## 3. Results

### 3.1. Model parameters produce oscillations in all frequency ranges

To ensure that our proposed model variations can produce oscillations in most physiological frequencies, we repeatedly simulated single node dynamics without any connectivity for 2-seconds while systematically varying the excitatory and inhibitory time constants. For each combination of the time constants, we examined whether the model produced an oscillatory wave form, and the peak frequency of the oscillations was extracted and assigned to a defined frequency band. Figure 2 clearly shows that the model is able to produce all frequencies up to 45 *Hz*. More importantly, the entire frequency range is covered by time constants ranging from 0 − 40 *ms*, which is consistent with most models [8, 49, 54, 24, 42]. For each frequency band, a characteristic waveform is shown with its corresponding power spectra. External input *P*(*t*) was set to *P*(*t*) = 2.5 to ensure the uncoupled model is in a limited cycle regime within the normal biological range for neuronal activity. The effect of the external drive parameter is shown in Figure S3, where the simulations show oscillatory behavior near *P*(*t*) = 2.5.

**Figure 2:**
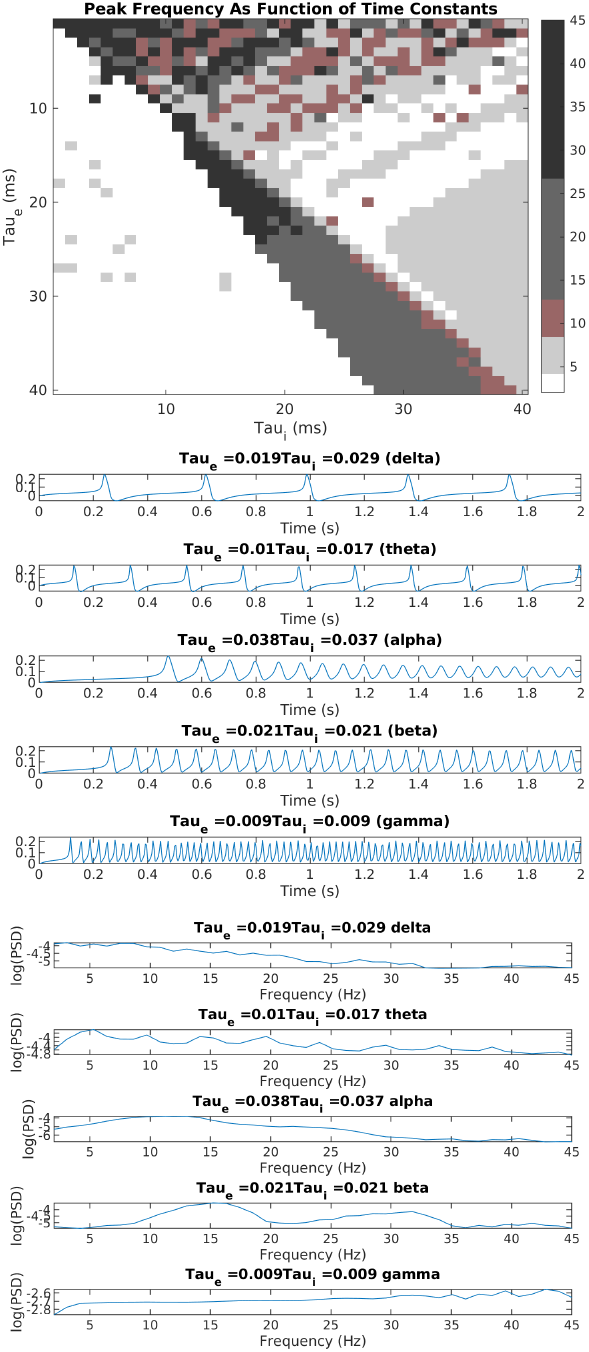
Peak frequency depends on time contants. Top: Heat map of models peak frequency (Hz) as a function of the excitatory and inhibitory time constants. Middle: oscillatory time course showing different peak frequencies, their corresponding power spectra is shown to the bottom.

**Figure 3:**
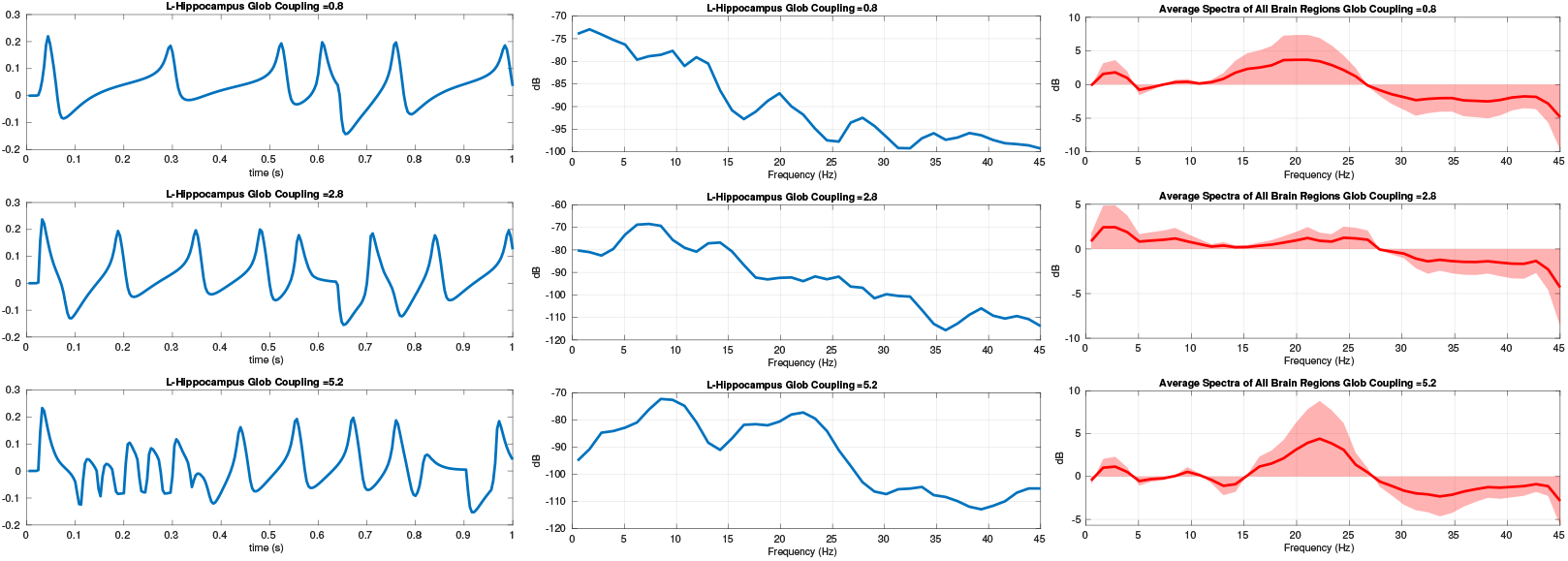
Global coupling controls oscillations. As the global coupling parameter increases, the simulated time series of a particular region is oscillating at higher frequencies as shown on the left column, each time series’ corresponding power spectra is shown in the middle. The right column shows the average spectra of all 86 brain regions after removing the mean. Transmission velocity between brain regions was held to a constant (10*m/s*) for all simulations.

**Figure 4:**
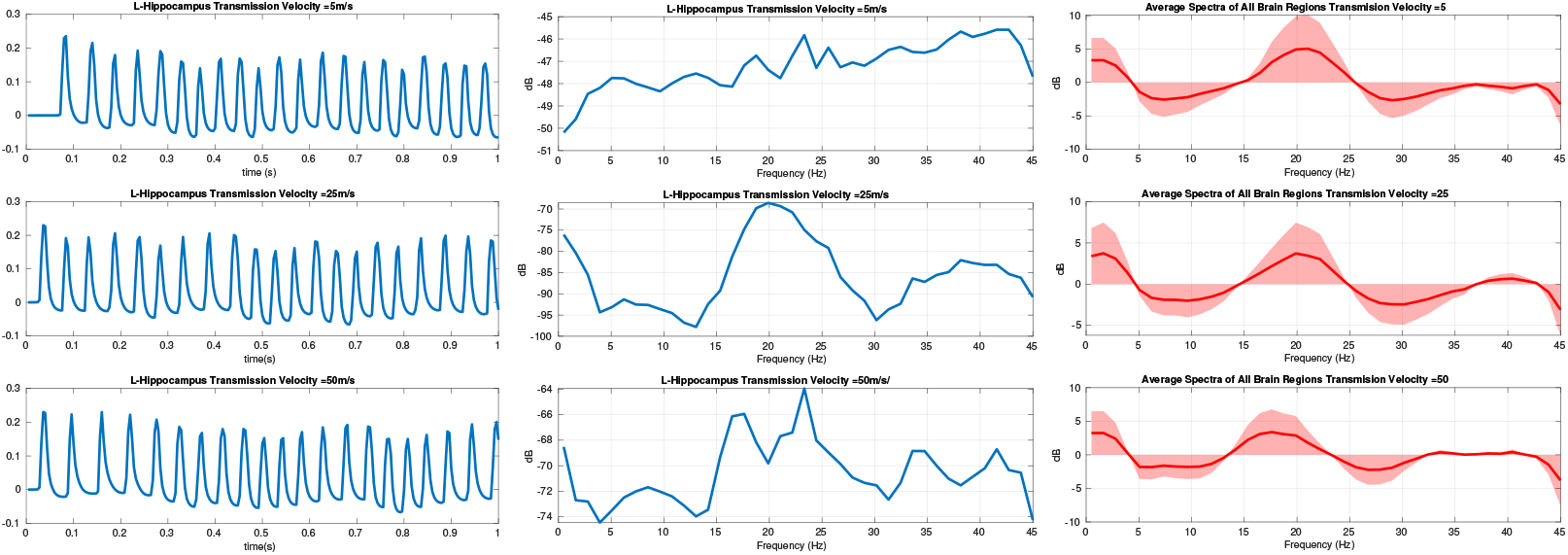
Transmission velocity and oscillatory behavior. In the network model, increasing the transmission velocity causes a time shift of the incoming signal; the left column shows the effect of the delay on 1 second simulated time course. The middle column shows the effect of transmission velocity on the corresponding power spectra. The right column shows the average spectra of all 86 brain regions after removing the mean. Global coupling was held to *c*_5_ = 1.5 for all simulations.

Using the same set of local parameters, we simulated the network dynamics of 86 interconnected regions using one structural connectivity matrix, a transmission velocity of ten meters per second, and varying global coupling parameter *c*_5_ ranging from 0 to 3. A representative subject’s structural connectivity i.e. weighted connectivity matrix whose elements represent the amount of fiber tracts connecting different regions, is given in Fig. S2, and the simulated time series and power spectra are shown in 3. The external input parameter was lowered to *P*(*t*) = 1.5 for these simulations to make sure that global coupling and connectivity was the main driver of oscillations (see Fig S3). The specific external input parameter value was chosen because Muldoon et al. [36] showed default model parameters injected with *P*(*t*) = 1.5 shifted the model from a low oscillatory state to a high oscillatory state. On the hand, using the same structural connectivity matrix, we simulated network dynamics while varying transmission velocity *v* ranging from 5 *m/s* to 50 *m/s* and holding global coupling to *c*_5_ = 1.5 for all simulation instances. 4 shows that transmission velocity has a subtle effect on the simulated time series. Despite changes in peak frequency for individual brain regions from slow speed (5 *m/s*) to faster speeds (25 *m/s* and 50 *m/s*), the average power spectrum from all simulated brain regions in the network remained the same regardless of transmission velocity *v*.

### 3.2. Optimized neural mass models

Most optimizations terminated upon reaching the maximum number of iterations allowed, which is *N_parameters_*× 500 iterations. However, the minimum within the boundary constraints was acquired before reaching the maximum iteration, the simulated annealing algorithm accepts additional function evaluations after acquiring a minimum to scan the rest of the parameter space, none of the optimization runs terminated while the cost-function evaluations were decreasing. None of the optimized parameters were reported to be equal to the upper or lower boundary, thus the specified range was not overly narrow, and a minimum was found within the bounds in all cases. The mean and standard deviation of all parameters are reported in Table 2. Recall that the three models we evaluated were: regionally varying oscillators (VO), regionally identical oscillators coupled by structural connectivity (IOC) and, regionally varying oscillators coupled by connectivity (VOC). We also evaluated the VOC model with iterative optimization of local and global parameters (denoted (VOC-CM). We observe that there is a difference between excitatory and inhibitory local parameters (*c*_1_*, c*_3_ and *c*_2_*, c*_4_ respectively), with the excitatory constants being consistently larger than inhibitory constants across all model variations. This slight variation between excitatory and inhibitory parameters in network models reflect physiological conditions and is crucial in producing functional neuronal activity. In terms of time constants, we see the excitatory term being slightly lower than the inhibitory term. Similarly, global coupling parameters are relatively low in VOC models compared to IOC, however, we see that IOC model parameters have high optimal values as well as high variation across all subjects, suggesting that higher connectome coupling is required to optimize the IOC model.

**Table 2:** Mean (standard deviation) of model parameters for all model variations. VO = Varying Oscillators, VOC = Varying Oscillators with connectivity, VOC-CM = Varying Oscillators with connectivity and optimized by CM, IOC = Identical Oscillators with connectivity.

|  | VO | VOC | VOC-CM | IOC |
| --- | --- | --- | --- | --- |
| Time constants (ms) | $\tau_e = 15.2(3.0)$<br>$\tau_i = 19.4(2.9)$ | | $\tau_e = 15.7(2.6)$<br>$\tau_i = 18.2(2.7)$ | $\tau_e = 18.1(9.0)$<br>$\tau_i = 24.8(8.8)$ |
| Local Parameters | $c_1 = 14.38(1.502)$<br>$c_2 = 9.989(2.166)$<br>$c_3 = 15.19(1.534)$<br>$c_4 = 6.117(1.794)$ | | $c_1 = 16.23$<br>$c_2 = 7.497(1.541)$<br>$c_3 = 16.63(0.955)$<br>$c_4 = 4.633(1.153)$ | $c_1 = 17.09(3.465)$<br>$c_2 = 5.032(3.743)$<br>$c_3 = 19.13(1.000)$<br>$c_4 = 4.082(2.711)$ |
| External Input | $P(t) = 2.664(0.094)$ | | $P(t) = 2.660(0.013)$ | $P(t) = 2.607(0.409)$ |
| Global Coupling | | $c_5 = 0.018(0.043)$ | $c_5 = 0.003(0.0075)$ | $c_5 = 5.093(3.697)$ |
| Transmission Velocity (m/s) | | $v = 8.714(4.455)$ | $v = 11.24(3.56)$ | $v = 11.75(5.506)$ |

**Table 3:** Table summarizing the Wilcoxon rank-sum test used to compare Kolmogorov-Smirnov statistics between different model implementations. All p-values reported were adjusted for multiple comparisons (Bonferoni).

|  | VO vs. VOC | VO vs. IOC | VOC vs. IOC | VO vs. VOC+CM | VOC vs. VOC+CM | IOC vs. VOC+CM |
| --- | --- | --- | --- | --- | --- | --- |
| Z-score | -27.21 | -27.78 | 2.149 | -11.85 | 26.29 | 27.98 |
| W statistic | $1.987 \times 10^5$ | $1.952 \times 10^5$ | $3.757 \times 10^5$ | $2.912 \times 10^5$ | $5.212 \times 10^5$ | $5.315 \times 10^5$ |
| P-Value | $P < 0.0001 **$ | $P < 0.0001 **$ | $P = 0.1899$ | $P < 0.0001 **$ | $P < 0.0001 **$ | $P < 0.0001 **$ |

Figure 5 (top) shows the cost-function values for the conditional minimization iterations over the global and local parameters in the VOC-CM optimization task. We see that the local parameter optimization iterations always result in a lower cost-function value than when optimizing over global parameters. However if we compare all of the global cost-function values and all the local cost-function values we see a downward trend in both that begins to flatten around iteration 7. Further iterations do not materially improve the fits, as it appears that the CM optimization has converged. The jaggedness of the curve also shows the importance of allowing an increase in the cost-function between the local- and global-steps, since otherwise no global step would improve upon the initial solution involving only local optimization. The CM performance for all all subjects is shown in Fig. S4.

**Figure 5:**
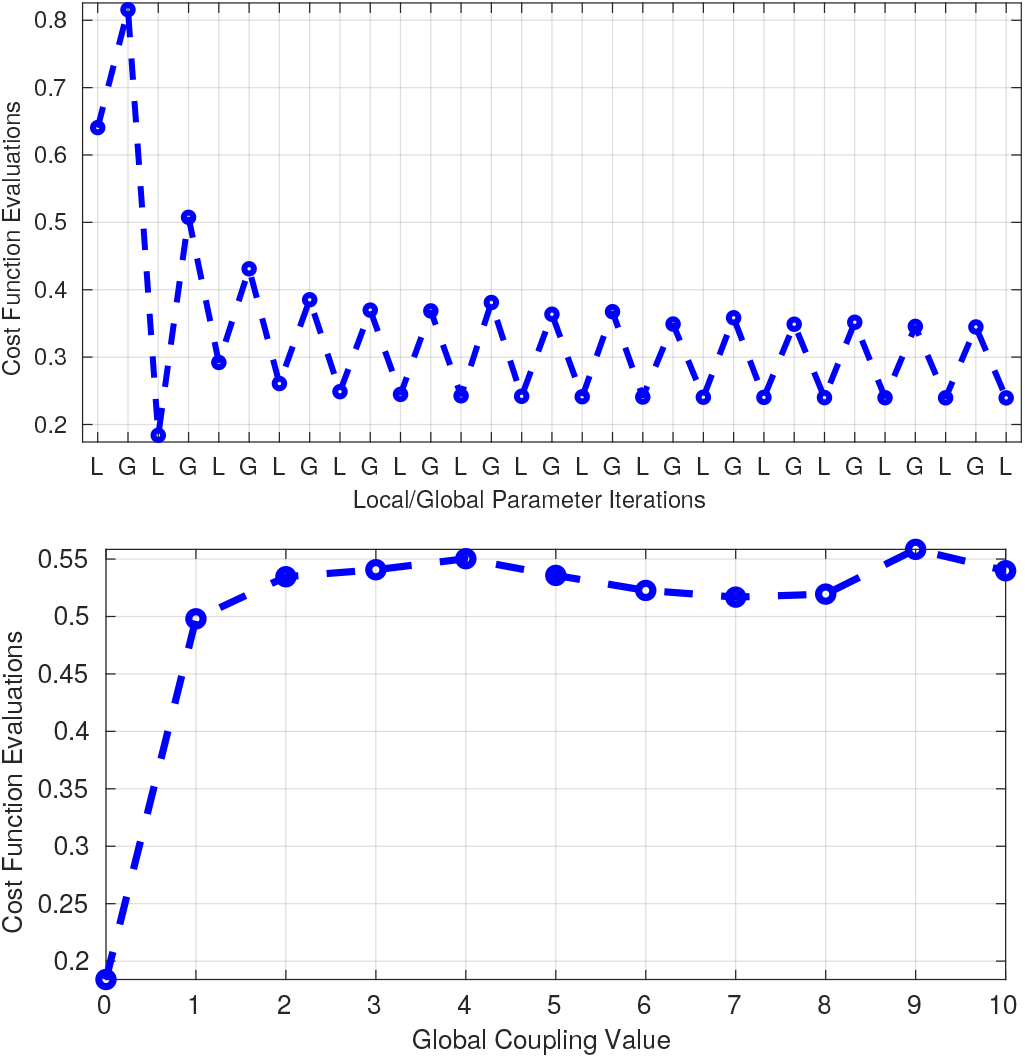
Top: Conditional minimization performance. The CM algorithm alternatively optimized local parameters and global parameters of the VCN model for 15 iterations. The optimized local parameters consistently resulted in lower cost-function evaluations than global parameters over all iterations. The final iteration was used as the set of optimized parameter for further analysis. Bottom: Global coupling parameter changes the parameter space. Introducing a structural connectivity matrix with increasing global coupling parameter increases the cost-function evaluation, but does not continuously increase the evaluations as global coupling increases.

To determine the effect of global coupling on model performance, we gradually increased the global coupling parameter in the VOC model while holding transmission velocity constant. We had hypothesized that introducing global coupling, structural connectivity, and transmission delay would improve the parameter space and yield a lower cost-function, but our results show the exact opposite. Figure 5 (bottom) shows that introducing global coupling is an uphill move in terms of cost-function evaluations and the corresponding changes in parameter space does not improve model performance. Alongside Fig 5 and 6, we see that re-optimizing for the global coupling and transmission velocity parameters in VOC cannot return the cost-function evaluations to the minimum achieved by local parameters only (VO).

Figure 6 (left) shows the boxplots of the Kolmogorov-Smirnov (KS) statistic between the source localized power spectra and its corresponding simulated power spectra from each model variation over each of the 86 brain regions in each of the 7 subjects. The best performing model was the individual oscillators fitted to the source localized spectra at each node (VO). VO and VOC-CM was able to minimize the KS-statistic by optimizing for each individual ROI, whereas IOC and VOC required minimizing for the average KS-statistic of all 86 ROIs, therefore a high variance around the median is shown in their box-plots. Contrary to our belief that connectivity improves fitting, introducing a connectome and global coupling to optimized oscillators resulted in higher cost-function evaluations (VOC). Using one set of local parameters for all brain regions in IOC produced similar results to VOC (*P* = 0.1899). On the other hand, optimizing the VOC model variation with the CM algorithm resulted in a much better model performance; the model fit of VOC-CM was significantly better than IOC and VOC (*P* < 0.0001).

**Figure 6:**
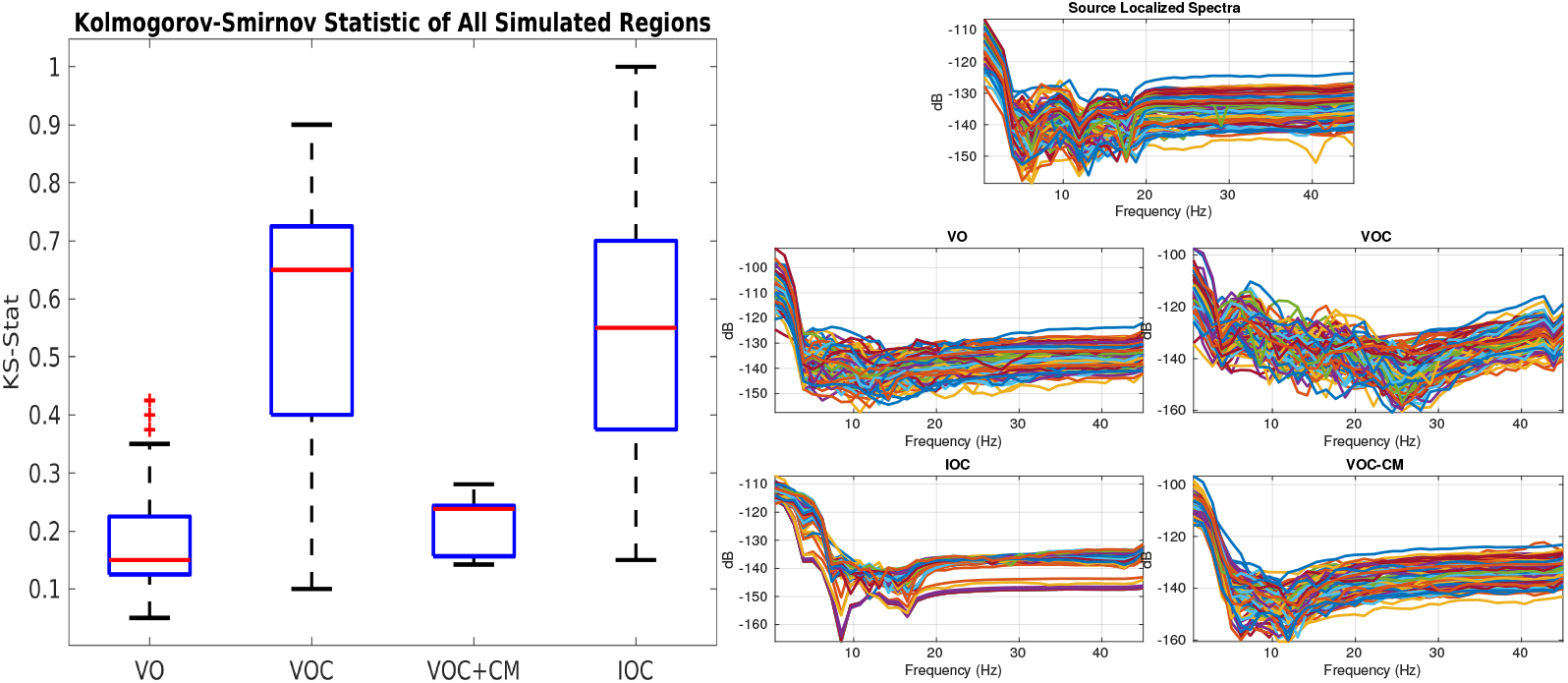
Comparison of model performance. Left: Summary of Kolmogorov-Smirnov statistics between different model variations (VO = varying oscillators, VOC = varying oscillators with connectivity, VOC + CM = varying oscillators with connectivity, optimized via CM, IOC = identical oscillators with connectivity) over all 86 ROIs and all 7 subjects. Right: Source localized power spectra and simulated model power spectra for all 86 regions averaged over all subjects.

The source localized power spectra of all regions and their corresponding simulated power spectra for each model variation are visualized in Fig. 6. The source localized spectra show a clear alpha peak at 8 − 12 *Hz* and a beta peak with lower power at near 20 *Hz*, which is characteristic of normal neurophysiological frequency profiles. Consistent with our KS-statistic results in Fig. 6, we see that the average IOC spectra does not show these characteristic peaks while other model variations do to a limited degree. The optimized parameters in Table 2 show relatively high variances in IOC compared to other models, and the parameter means between excitatory and inhibitory time constants differ by a small amount, suggesting the optimization algorithm had trouble converging onto a parameter range that is suitable for this model variation. The consequence of having identical parameters for each node and small differences between excitatory and inhibitory parameters for IOC is shown in Fig. 7, where each region’s spectra are less likely to have various peaks and troughs. Despite the VO and VOC-CM spectra having a lower KS-statistic than other spectra in Fig. 6, their beta activity is not as distinct as what’s shown in the source localized spectra. Finally, with the exception of IOC, the remaining model variations recapitulates the observed alpha peaks in the source localized spectra to a limited degree.

**Figure 7:**
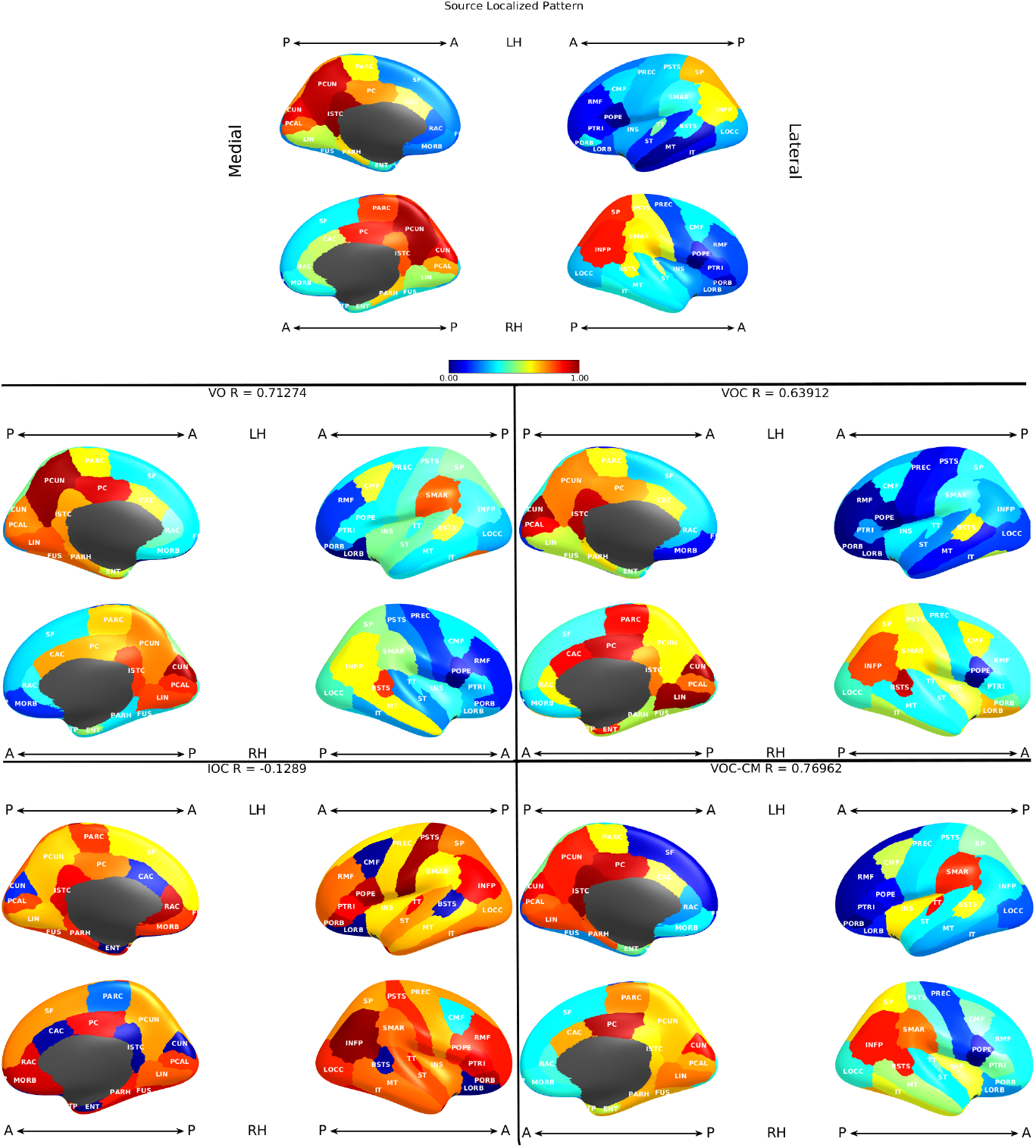
Spatial distribution of alpha band. Surface plots showing the power in the alpha band averaged across all subjects. From top to bottom: (1) empirical data, (2) varying oscillators (VO) model, (3) varying oscillators with connectivity (VOC), (4) identical oscillators with connectivity (IOC), and (5) varying oscillators with connectivity, optimized using conditional minimization (VOC-CM). The color intensity indicates the amount of power within the 8 − 12*Hz* range in the frequency domain, scaled by the mean of the alpha power over each region.

### 3.3. Spatially distributed patterns of power spectra

Figure 7 illustrates via surface-plots the alpha band power (8 − 12 *Hz*) over the entire brain for the observed and simulated spectra averaged from all subjects. Each of the cortical regions are colored by the intensity of that region’s alpha power scaled by the mean alpha power over all brain regions. As expected, the source localized spectra (top row) shows relatively larger spheres in the posterior regions of the brain. The VO, VOC, and VOC-CM models show the same trend, although they are distributed more laterally than the observed alpha distribution. The IOC model did not match the alpha spectra spatial pattern at all, with only a small number of regions that contain alpha powers significantly above the mean. The Pearson’s correlation coefficients are displayed on top of each glass-brain plot, and as expected, VO and VOC-CM had the highest correlation when comparing the 86 brain region’s alpha powers.

From the optimization results in Figure 5, we already see a change of less than 1% in cost function evaluations as the conditional minimization algorithm approached the 10*^th^* iteration, suggesting any of the solutions along the end of the conditional minimization algorithm could be a plausible solution. We selected parameter sets that computed cost-function evaluations within ± 1% range of the final cost-function evaluation. The probability distribution of these optimized parameters are shown in Figure 8. The majority of the parameters from varying oscillators (VO) model shows a bimodal distribution, with many peaks in the histogram suggesting different viable solutions that satisfies our goodness-of-fit criteria. On the other hand, the parameters chosen from the final iteration of the VOC-CM model shows a less obvious bimodal distribution with the exception of *τ_i_*. Additionally, the histogram peaks suggest that there are at least two highly probable parameter values for each parameter in both cases. Despite conditional minization converging to a low cost-function evaluation that drop less than 1% after the 10th iteration, the parameters were still unable to converge to a single value, further emphasizing the difficulty of finding unique solutions to an over-specified model.

**Figure 8:**
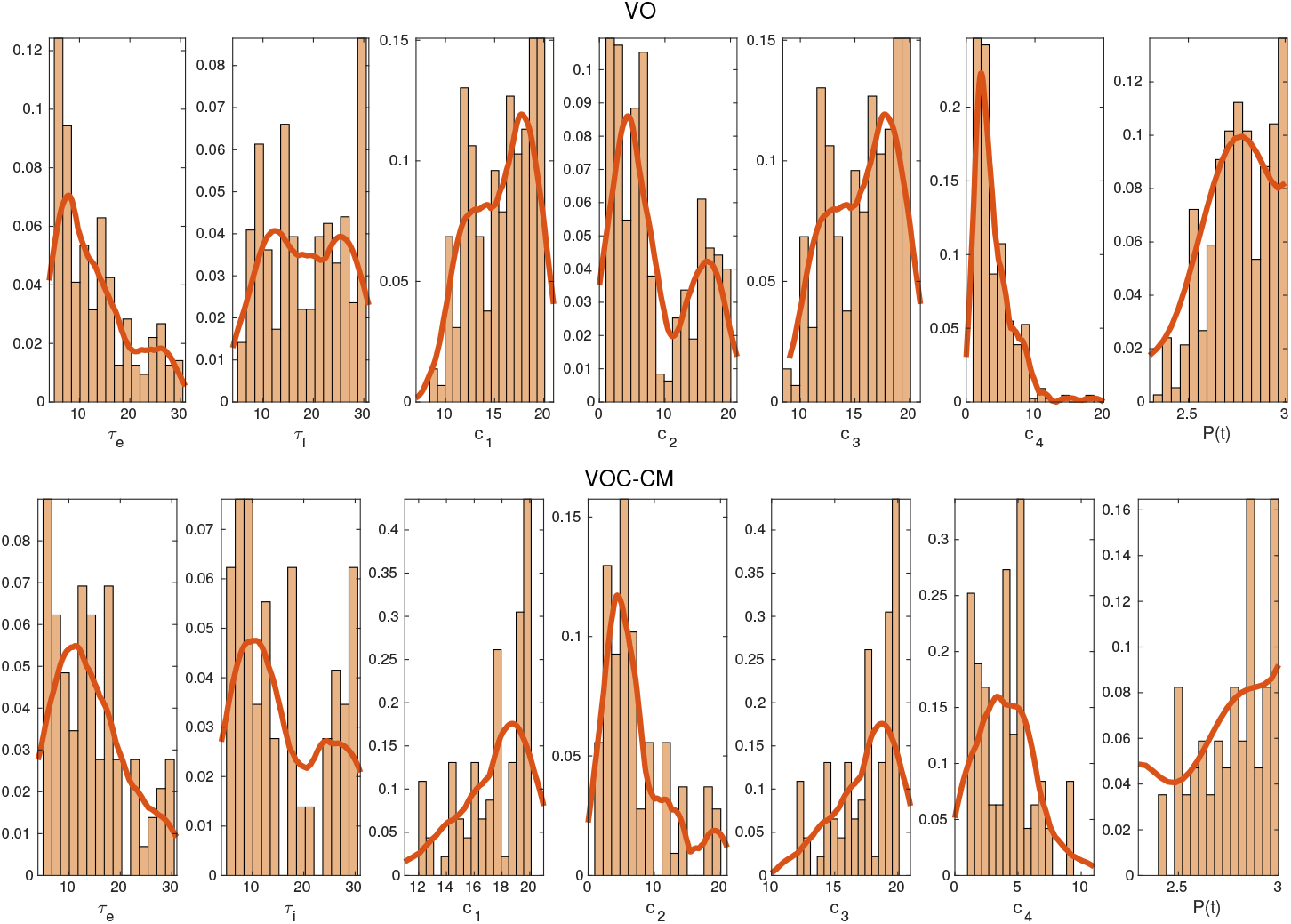
Best Fitting Model Parameters. Histograms showing the probability distribution of parameters chosen from ± 1% of the best fitting solution for the varying oscillators (VO, top) model and varying oscillators with connectivity, optimized using conditional minimization (VOC-CM, bottom).

## 4. Discussion

A challenge for emerging models of brain activity is that in a complex dynamical system such as the brain, it is difficult to predict function even if the underlying architecture, local cortical dynamics, and cortical-cortical interactions are known. In the present article, we studied the role of local and global parameters in a system of coupled oscillating neural mass (Wilson-Cowan) models, either unconnected or connected via white matter fibers as measured from diffusion-MRI. As described in previous network modeling efforts, coupled dynamical systems have a collective behavior that depends on the network structure, the local dynamics of each node, and the coupling function for the transfer of information [45, 21, 34]. Using different implementations of the Wilson-Cowan oscillator model, we reproduced to varying degrees of success spatially varying spectral features of human source localized EEG at rest. Our results show that 1) introduction of the connectome to the oscillator model does not improve model fitting to source localized EEG, 2) the identifiability problem manifests itself in the model’s parameter space as well as the spatial distribution of the modeled frequency profile.

First, we aimed to determine which configuration of our chosen neural mass model best reproduces source localized EEG data. From our simulations, it is clear that a model of individual oscillators at each brain region (VO) is capable of reproducing the spatial and spectral patterns of EEG data. While the absence of network topology in the VO model does not correctly depict the interconnected brain regions, the one pair of oscillator model per brain region fitting criteria is a much easier parameter inference problem than inferring network model parameters. VOs simulations produced a mean KS-stat of around 0.15, which is lowest out of all model variations. On the other hand, network models (VOC, VOC-CM) were also able to produce the alpha and beta spatial patterns that closely matched our source localized EEG data. The Jansen and Rit model [24] utilized realistic ratios of excitatory and inhibitory connections in a neuronal ensemble to arrive at their parameter values, David and Friston [8] expanded on this idea and established a neural mass model with similar differences between excitatory and inhibitory parameters. Interestingly, IOC parameters exhibiting this difference between excitatory and inhibitory parameter values were not able to produce a satisfactory spectra or a posteriorly distributed alpha pattern, indicating the importance of allowing spatially varying local parameters in order to produce characteristic neuronal patterns. In the IOC model, the only terms driving regional differences in the brain were the connectivity matrix, global coupling, and tramission velocity, which is an indirect way of determining the effect of introducing a connectome to an optimized network. Surprisingly, despite it’s anatomical relevance, the structural connectivity does not improve the model performance, but drastically alters the parameter space instead.

Our results show a simple addition of network connectivity to individual oscillators optimized independently at all brain regions does not improve the performance of the model. As shown in Figure 5 and 6, no amount of connectome coupling, while keeping the VO local parameters, improves model performance; in fact, it makes is substantially worse, with the KS-statistic cost function plateauing around 0.5-0.55 as global coupling increased gradually, compared to the KS cost of the VO model of less than 0.2. We conjecture that the one-to-one fitting without any connectivity and transmission velocity influences may have provided a simpler optimization problem than the network models. Because we used optimized local parameters from the VO model in VOC model, we expected similar or better model performance with the addition of a more physiological, interconnected brain network. However, despite optimized local dynamics at each node, the interconnected regions introduced an uphill move for the optimization algorithm instead of a downhill move, suggesting the feedback from adjacent regions may be changing local dynamics that are not explainable by just a global coupling parameter and transmission velocity. Our conditional minimization algorithm was able to optimize our local and global parameters iteratively until we obtained a set of parameters that outperformed VOC. As described above, despite IOC having identical nodes, meaning only one set of local parameters for the entire network, the inferred parameters are high in variance and do not reflect neurophysiological conditions. This is consistent with the findings by [2], suggesting that network dynamics do not only depend on anatomical connectivity, but also on “state-dependent dynamical regimes of the brain regions” and on the heterogeneity of node degrees.

The surface-plots displaying the spatial distribution of each model variation’s alpha pattern highlight the identifiability problem of network neural mass models. Despite the differences in parameterization, all models show spatial alpha patterns that are identical to each other with the exception of IOC. In the frequency domain, there are recognizable differences in the power spectra produced by each model, however, the minor differences do not necessarily capture the neurophysiological oscillations that translates to function. Additionally, the histograms in 8 shows there are many probable solutions that provide satisfactory spectra according to our goodness-of-fit criterion.

To capture function deteriorations in a diseased brain by mathematical models, there has been many recent attemps to correlate neural mass model parameters with stroke recovery [13, 12, 1], Alzheimer’s disease [58], and epilepsy [27]. However, all these efforts neglect the over-parameterization of the models by expanding neural masses to networks in order to maximize a fit to functional connectivity. Correlating a set of parameters with a change in functional connectivity does not mean such parameter shifts are meaningful enough to diagnose disease, as another set of parameters may capture the same functional connectivity just as well. Our results show the manifestation of identifiability problems in neural mass models as a challenge to diagnosing disease via mathematical models, as network models need to capture both functional and spatial information in order to fully capture disease spread. During parameter inference, careful inspection of the parameter distribution and model behavior is needed to obtain parameters that converged to a uniform distribution. We believe low dimensional models with parameter constraints may avoid the identifiability problem and provide more meaningful model parameters.

## 6. Supplementary Material

**Figure S1:**
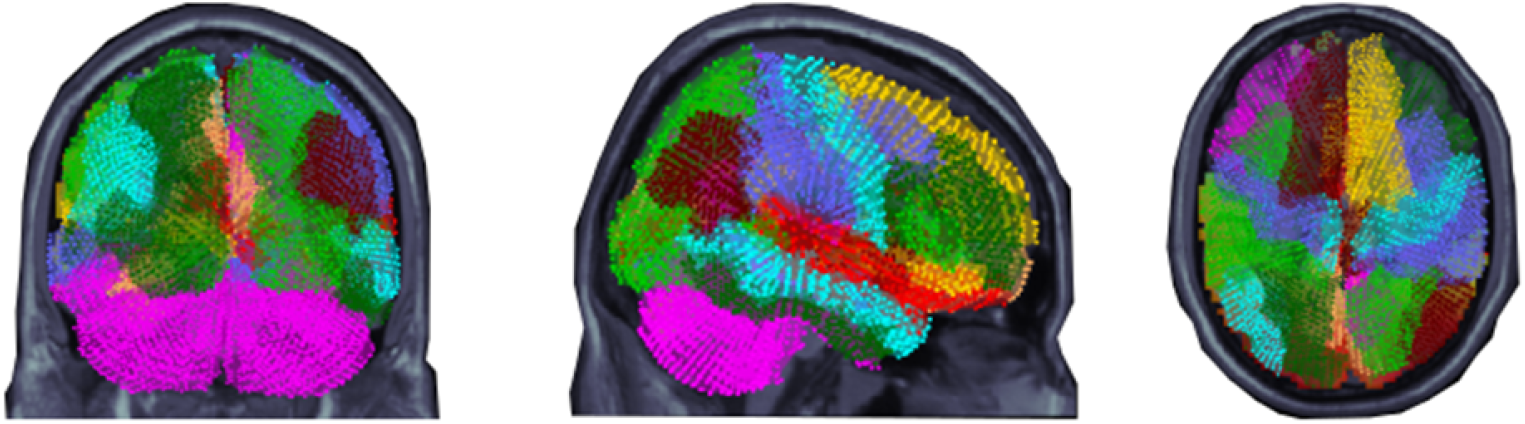
Dots representing volumetric source locations mapped to their respective regions of interest (ROI) viewed from the back, right, and top. Different colors represent the 86 segmented regions in the FreeSurfer Desikan-Killany atlas, each ROI is viewed as a node on the connectome.

**Figure S2:**
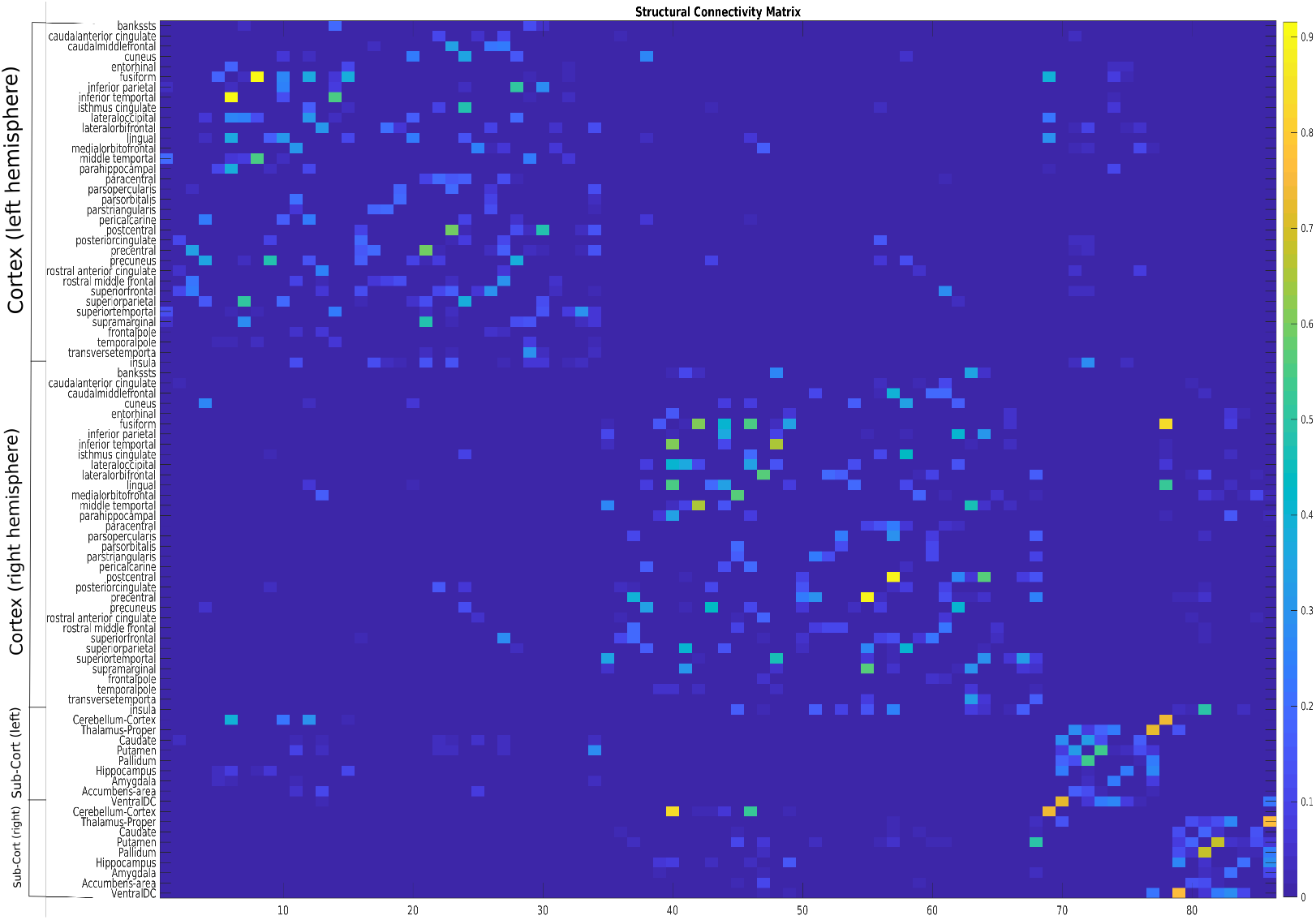
Structural connectivity matrix of one representative subject.

**Figure S3:**
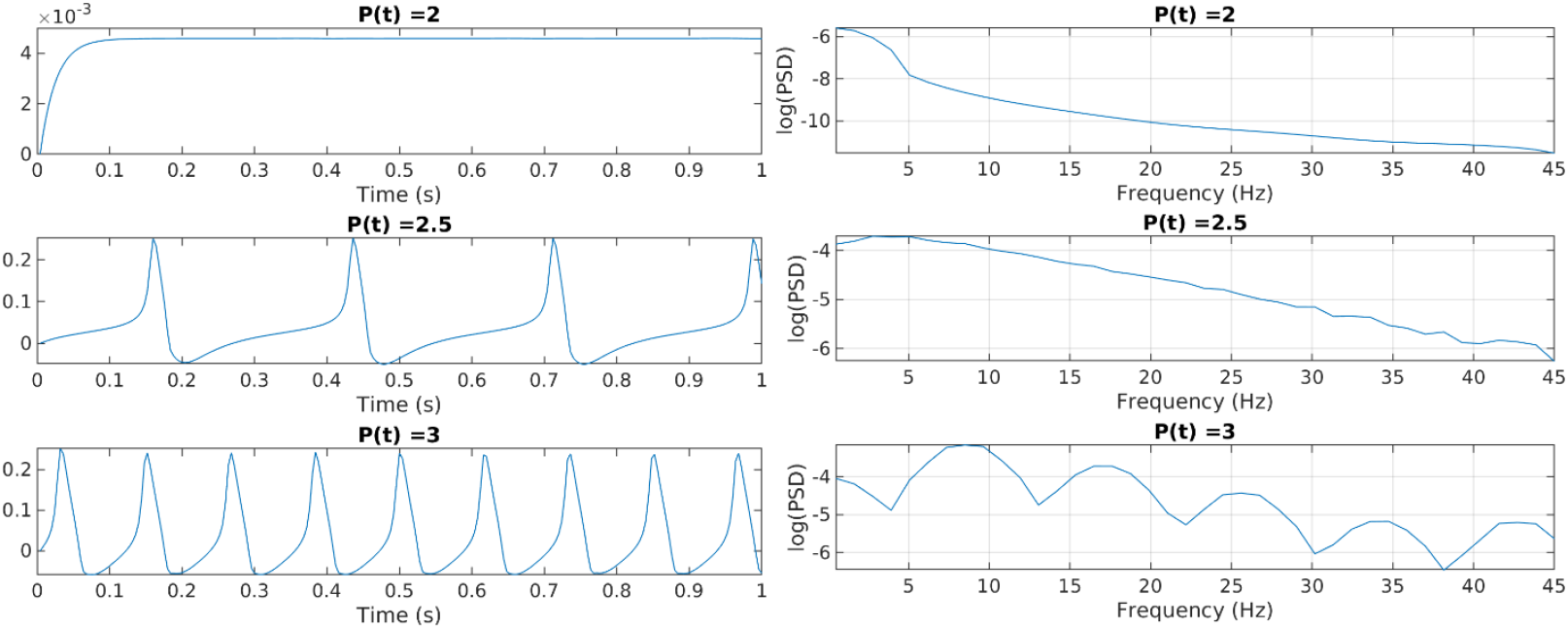
Neural mass model’s oscillatory activity changes as external drive parameter *P*(*t*) is gradually increased at one node.

**Figure S4:**
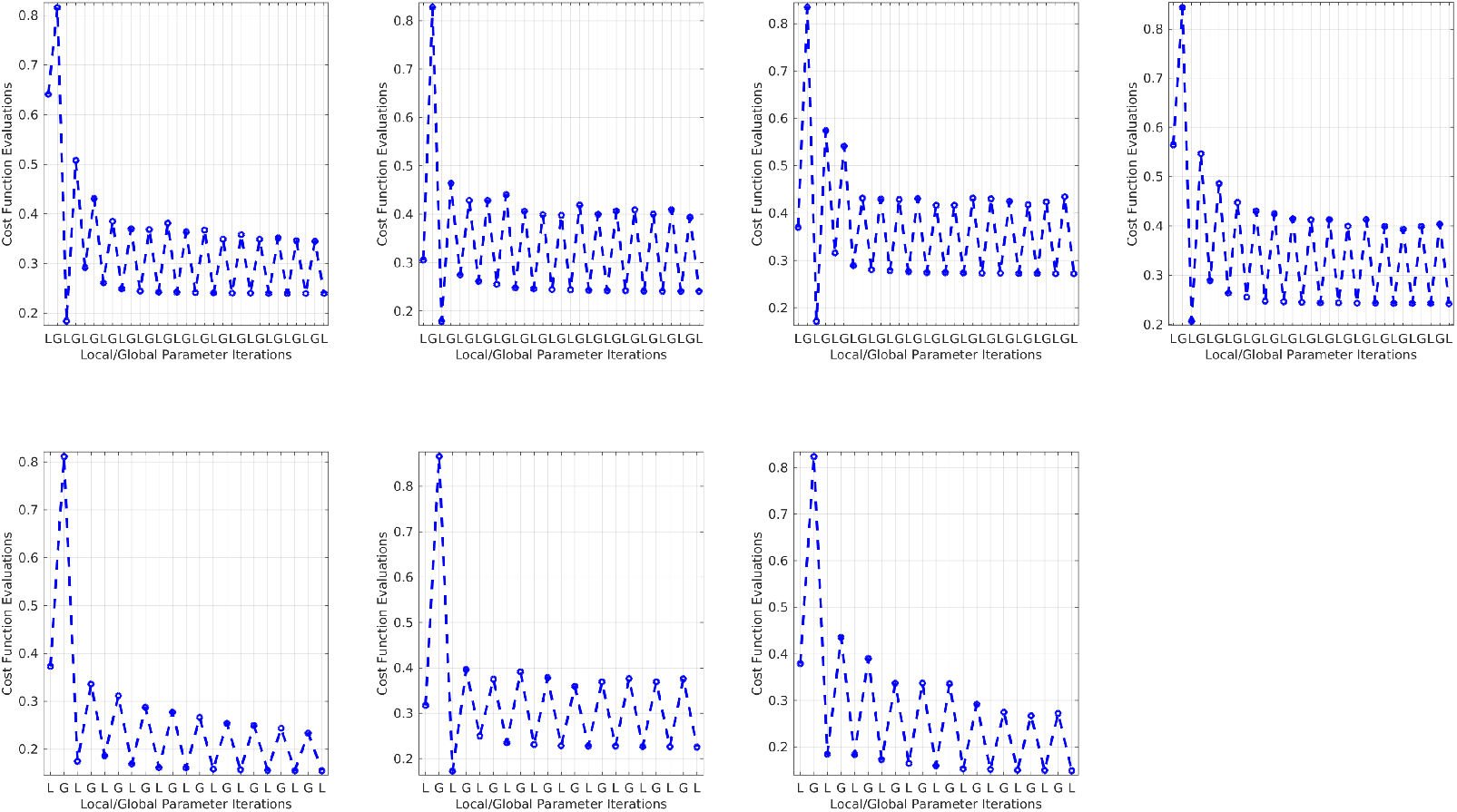
Conditional minimization performance for all subjects.

